# Senescence of endplate osteoclasts induces sensory innervation and spinal pain

**DOI:** 10.1101/2023.10.26.564218

**Authors:** Dayu Pan, Kheiria Gamal Benkato, Xuequan Han, Jinjian Zheng, Vijay Kumar, Mei Wan, Junying Zheng, Xu Cao

## Abstract

Spinal pain affects individuals of all ages and is the most common musculoskeletal problem globally. Its clinical management remains a challenge as the underlying mechanisms leading to it are still unclear. Here, we report that significantly increased numbers of senescent osteoclasts (SnOCs) are observed in mouse models of spinal hypersensitivity, like lumbar spine instability (LSI) or aging, compared to controls. The larger population of SnOCs is associated with induced sensory nerve innervation, as well as the growth of H-type vessels, in the porous endplate. We show that deletion of senescent cells by administration of the senolytic drug Navitoclax (ABT263) results in significantly less spinal hypersensitivity, spinal degeneration, porosity of the endplate, sensory nerve innervation and H-type vessel growth in the endplate. We also show that there is significantly increased SnOC-mediated secretion of Netrin-1 and NGF, two well-established sensory nerve growth factors, compared to non-senescent OCs. These findings suggest that pharmacological elimination of SnOCs may be a potent therapy to treat spinal pain.

## Introduction

Low back pain (LBP) is the most common musculoskeletal problem globally, affecting at least 80% of all individuals at some point in their lifetime^1–4^. It has also become the leading cause of years lived with disability (YLDs) worldwide, with almost 65 million cases involved per year^5,6^, which consequently results in a tremendous medical burden and economic cost ^2,7^. Current pharmacologic treatment options for LBP include nonsteroidal anti-inflammatory drugs (NSAIDs), corticosteroids, opioids, etc^8^. However, the clinical use of these medications is limited due to potential severe adverse effects and modest therapeutic efficacy^9–11^. Some biological agents aiming for LBP management are under investigation as well^12,13^. For example, parathyroid hormone (PTH) has shown a superior antinociceptive effect on LBP in recent studies^14,15^, whereas its risk of causing osteosarcoma and Paget’s disease is not negligible^16–18^. Thus, there is an urgent unmet clinical need for effective nonsurgical therapeutic interventions for LBP.

Cellular senescence is a stable and terminal state of growth arrest in which cells are unable to proliferate despite optimal growth conditions and mitogenic stimuli^19^. It can be induced by various triggers, including DNA damage, telomere dysfunction and organelle stress, and it has been linked to host physiological processes and age-related diseases, such as atherosclerosis, type 2 diabetes and glaucoma^20–23^. Hence, the clearance of senescent cells has been suggested as a promising therapeutic strategy in several areas of pathology. For instance, glomerulosclerosis and decline in renal function in aged mice are rescued by clearance of p16^INK4a^-expressing senescent tubular brush-border epithelial cells, while clearance of senescent cells reduces age-related cardiomyocyte hypertrophy and improves cardiac stress tolerance^24^.

Likewise, it has been shown that cellular senescence is an essential factor in the promotion of age-related musculoskeletal diseases, such as osteoporosis^25^, osteoarthritis^26^ and intervertebral disc (IVD) degeneration^27,28^. The effectiveness of senolytic drugs towards bone related diseases via elimination of senescent cells is also well-documented^25^. In particular, for the treatment of IVD degeneration, which is strongly associated with LBP, it was demonstrated that the senescent cells of degenerative discs were removed, and the IVD structure was restored by treatment with ABT263, a potent senolytic agent, in an injury-induced IVD degeneration rat model^29^. Furthermore, recent research shows a direct link between telomere shortening-induced cellular senescence and chronic pain hypersensitivity^30^. Previous studies conducted by our laboratory have elucidated the significant role of osteoclasts in initiating the porosity of endplates with sensory innervation into porous areas and triggering LBP^31,32^. Attenuating sensory innervation by inhibiting osteoclast activity could reduce spinal pain sensitivity. Importantly, it has been observed that some osteoclasts exhibit characteristics of senescence during osteoclastogeneses, such as the expression of p16 and p21, appearing to obtain a heterogeneous senescent phenotype^33–35^. Based on all these findings, we speculated there might be a subgroup of osteoclasts that are senescent (which we refer to as SnOCs), and they might promote the induction of sensory nerve innervation in porous endplates and spinal hypersensitivity.

Here, we demonstrate the presence of SnOCs in the porous endplates in two different spinal hypersensitivity mouse models induced by aging and LSI, respectively. We then deleted SnOCs in these models with ABT263, which resulted in a decreased number of tartrate-resistant acid phosphatase positive (TRAP^+^) OCs in the endplates along with decreased endplate porosity, reduced sensory innervation and attenuated spinal pain behaviors. Together, these findings suggest the potential of utilizing senolytic drugs for the treatment of LBP and its associated pathologies.

## Results

### A significantly increased number of SnOCs are associated with endplate degeneration and spinal hypersensitivity in the LSI and aged mouse models

In this study, we used two different LBP mouse models created by LSI and aging. LSI was induced in 3-month-old C57/BL6 mice by surgically resecting the L3–L5 spinous processes along with the supraspinous and interspinous ligaments^36–38^. Aged (24-month-old) C57BL/6J male mice were purchased from Jackson Laboratory. To explore spinal hypersensitivity, pain-related behavioral assessments, such as the von Frey test, hot plate test and active wheel test, were performed on sham-operated mice, LSI mice and aged mice. In both models, there was significantly less active time, distance traveled, and maximum speeds compared to the sham control (Figure 1a-c). Additionally, LSI mice and aged mice displayed significantly less reduced heat response times (Figure 1d), as well as significantly increased frequencies of paw withdrawal (PWF), depending on the strength of the mechanical stimulation, (Figure 1e, 1f) compared to the sham mice. By three-dimensional microcomputed tomography (μCT) analysis we found a significant increase in the porosity and separation of trabecular bone (Tb.Sp) within the endplates of both LSI and aged mice compared to their younger counterparts without LSI (Figure 1g, 1k).

**Figure 1.**
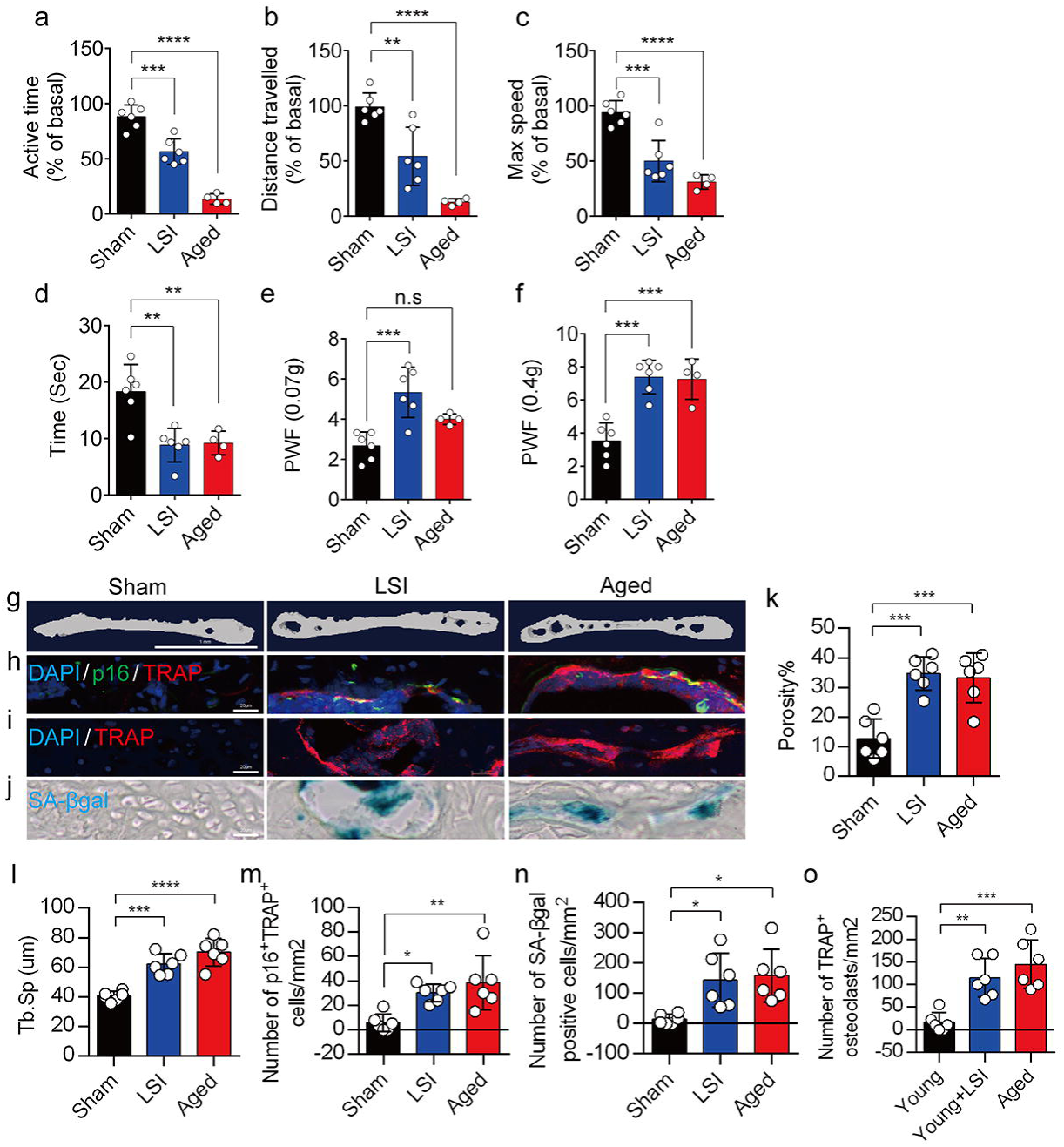
A greater number of SnOCs are associated with endplate degeneration and spinal hypersensitivity in the LSI and aged mouse models. (**a-c**) Spontaneous activity, including active time (**a**), distance traveled (**b**), and maximum speed (**c**) on the wheel within 48 hours in the sham, LSI-injury and aged mice. (**d**) Time in seconds spent on a hot plate in the three groups of mice. (**e, f**) The frequency of hind paw withdrawal (PWF) in response to mechanical stimulation (von Frey test, 0.07g (**e**) and 0.4g (**f**) in the sham, LSI-injury and aged mice. (**g**) μCT images of coronal caudal endplate sections of L4-5 from 3-month-old sham and LSI- and 24-month-old aged mice. (**h**) Immunofluorescent (IF) staining of p16 (green), TRAP (red), and DAPI (blue) of the endplates of sham, LSI-, and aged mice. (**i** and **j**) IF staining of TRAP (red) and DAPI (blue) (**i**) and SA-β-gal (blue) staining (**j**) of endplate serial sections of sham, LSI-surgery, and aged mice. (**k** and **l**) μCT quantitative analysis of the porosity percentage (**k**) and trabecular separation (Tb. Sp) (**l**) of the endplates in the indicated groups. (**m**) Number of SnOCs (p16-positive and TRAP-positive cells) per mm^2^ in the indicated groups. (**n**) Number of SA-β-gal (blue) positive cells per mm^2^ in the endplates in the indicated groups. (**o**) Number of TRAP (red) positive cells per mm^2^ in the endplates in the indicated groups. *n* ≥ 4 per group. Scale bar, 1 mm (**g**) and 20 μm (**h, i, j**). Statistical significance was determined by one-way ANOVA, and all data are shown as means ± standard deviations.

To investigate the potential relationship between SnOCs and the degeneration of spinal endplates in the context of LSI and aging, co-staining of TRAP (tartrate-resistant acid phosphatase), a glycosylated monomeric metalloprotein enzyme expressed in osteoclasts, and p16, a tumor suppressor and established marker for cellular senescence, was performed (Figure 1h). We found that compared to the control young sham mice, there was a significantly increased number of p16^+^TRAP^+^ cells in the LSI and aged mice, indicative of SnOCs occurring in the endplates of these two mouse models (Figure 1h, 1m). To confirm the occurrence of SnOCs in the endplates in the LSI and aged mice, we stained the adjacent slides with TRAP and Senescence-associated beta-galactosidase (SA-βGal) or HMGB1, markers for cellular senescence, respectively, and found that SnOCs existed in the two LBP mouse models but not in the sham controls (Figure 1i, 1j, 1n, 1o, Figure 1-figure supplement 1a, 1b).

These findings collectively show a strong association between the presence of SnOCs and the development of spinal hypersensitivity, along with the degenerative changes in the endplates of mice subjected to LSI and during the aging process.

### ABT263 effectively depletes endplate SnOCs in the LSI and aging mouse models

To study the contribution of SnOCs to spinal pain, we first needed to show that such cells could be successfully depleted. Thus, we treated 24-month-old aged mice and 3-month-old sham and LSI mice with ABT263, a specific inhibitor targeting the anti-apoptotic proteins BCL-2 and BCL-xL, effectively leading to the depletion of SnOCs^39^. ABT263 was administered via gavage at a dose of 50 mg/kg per day for seven days per cycle, with two cycles separated by a 2-week interval, resulting in a total treatment period of 4 weeks. Remarkably, after the administration of ABT263, we observed a significant reduction of SnOCs in the endplates compared to the PBS-treated group (Figure 2a-c).

**Figure 2.**
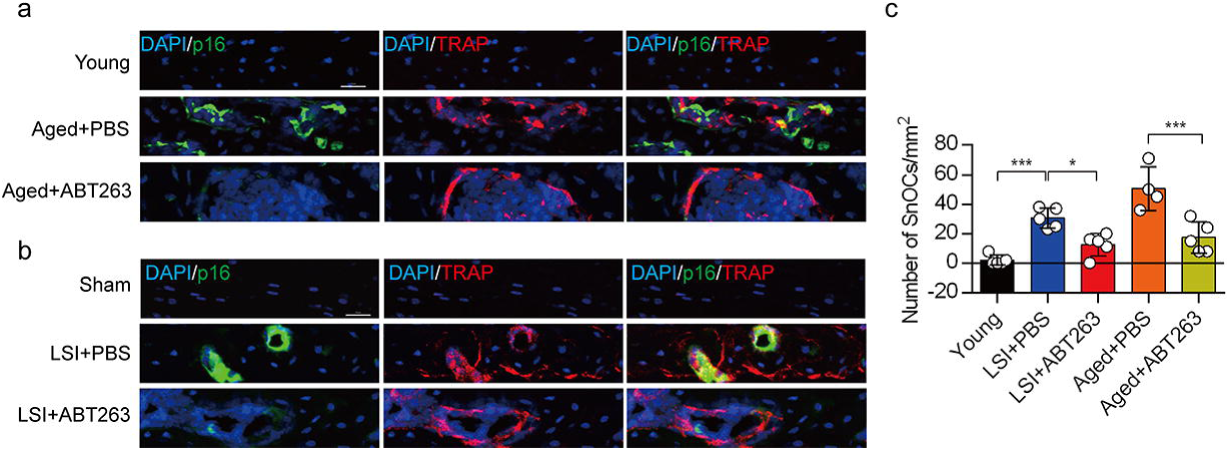
ABT263 effectively depletes endplate SnOCs in the LSI and aging mouse models. (**a-c)** Immunofluorescent staining of p16 (green), TRAP (red) and nuclei (DAPI; blue) of the endplates in aged (**a**) and LSI-mice (**b**) injected with PBS (control) or ABT263 and the quantitative analysis of SnOCs based on dual staining for p16 and TRAP (**c**). *n* ≥ 4 per group. Scale bar, 20 μm. Statistical significance was determined by one-way ANOVA, and all data are shown as means ± standard deviations.

### Eliminating SnOCs reduces spinal hypersensitivity

To investigate spinal hypersensitivity, pain behavioral tests, including the von Frey test, hot plate test and active wheel test, were conducted in sham, LSI- and aged mice treated with ABT263 and PBS, respectively. In the aged mice, ABT263 treatment resulted in a significant reduction in PWF (Figure 3a, 3b) and prolonged heat response times (Figure 3c) compared to the PBS-treated mice. In addition, there is a significant increased paw withdrawal frequency (PWF) in aged mice treated with PBS compared with young mice, particularly at 0.4g instead of 0.07g (Figure 3a, 3b). Furthermore, aged mice treated with PBS exhibited a significant reduction in both distance traveled and active time when compared to young mice (Figure 3d, 3e). Additionally, PBS-treated aged mice demonstrated a significantly shortened heat response time relative to young mice (Figure 3c). Importantly, these aged mice treated with ABT263 exhibited significantly increased distance traveled and active time compared to aged mice that received PBS injections (Figure 3d, e).

**Figure 3.**
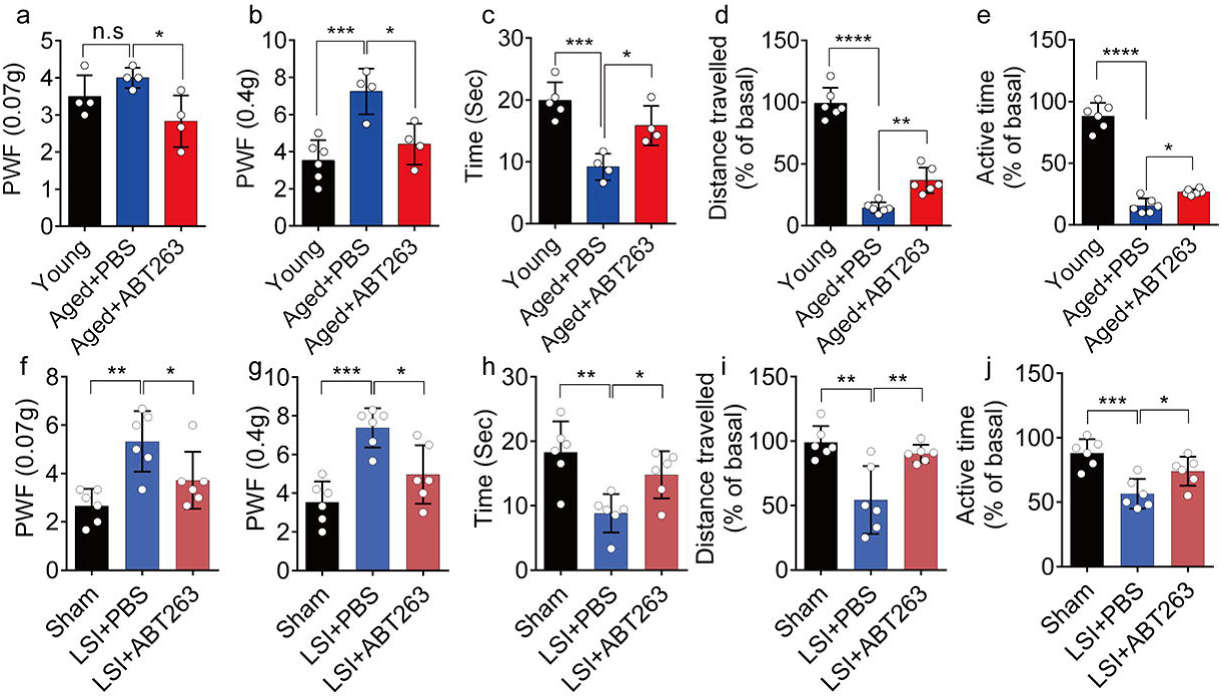
ABT263 treatment improves the symptomatic spinal pain behavior in the aged and LSI mouse models. (**a, b**) The PWF in response to mechanical stimulation (von Frey test, 0. 07g (**a**) and 0.4g (**b**) in aged mice treated with PBS or ABT263 compared to young adult mice. (**c-e**) Time (in seconds) spent on a hot plate (**c**), as well as spontaneous activity, including distance traveled (**d**) and active time (**e**), on the wheel within 48 hours in aged mice treated with PBS or ABT263 compared to young adult mice. (**f, g**) The PWF in response to mechanical stimulation (von Frey test, 0. 07g (**f**) and 0.4g (**g**)) in the LSI mouse model treated with PBS or ABT263 compared to sham-operated mice. (**h-j**) Time (in seconds) spent on a hot plate (**h**), as well as spontaneous activity analysis, including distance traveled (**i**) and active time (**j**) on the wheel within 48 hours in the sham and LSI-mice treated with PBS or ABT263. *n* ≥ 4 per group. Statistical significance was determined by one-way ANOVA, and all data are shown as means ± standard deviations.

Notably, LSI mice treated with ABT263 also demonstrated substantial improvements across several parameters compared to the PBS-treated control mice. These improvements included lower PWF (Figure 3f, 3g), prolonged heat response time (Figure 3h), increased distance traveled (Figure 3i), and extended active time (Figure 3j). These results collectively indicate that the elimination of SnOCs reduces spinal hypersensitivity in both aged and LSI mouse models.

### Depletion of SnOCs reduces spinal degeneration and sustains endplate microarchitecture

To determine the effect of SnOCs on endplate architecture, degeneration, and osteoclast formation in the context of spinal pain in the aged mice, we conducted μCT analysis and immunostaining in aged mice treated with ABT263, or PBS and untreated 3-month-old young mice (Figure 4a-e). There was a significant reduction in endplate porosity and trabecular separation (Tb.Sp) of the caudal endplates of L4/5 in the aged mice treated with ABT263 compared to the PBS-treated aged group (Figure 4f, 4g). The PBS-treated aged mice exhibited a significant increase in endplate porosity (Figure 4f) and trabecular separation (Tb.Sp) (Figure. 4g) compared to young mice. To examine the effects of ABT263 on endplate degeneration, we performed Safranin O staining and immunofluorescent staining to target matrix metalloproteinase 13-containing (MMP13^+^) and type X collagen-containing (ColX^+^) components within the endplate (Figure 4b-d). In the aged model, the ABT263-treated group exhibited a significant reduction in endplate score (Figure 4h), as well as the distribution of MMP13 and ColX within the endplates, compared to aged mice treated with PBS (Figure 4i, 4j). PBS-treated aged mice showed a significant elevation in endplate score (Figure 4h), as well as an increased distribution of MMP13 and ColX within the endplates when compared to young mice (Figure. 4i, 4j). Furthermore, TRAP staining demonstrated a substantial rise in the count of TRAP^+^ osteoclasts within the endplates of aged mice in comparison to young control mice. Importantly, ABT263 treatment resulted in a significant reduction of TRAP^+^ osteoclasts within the endplates compared to PBS treatment (Figure 4e, 4k).

**Figure 4.**
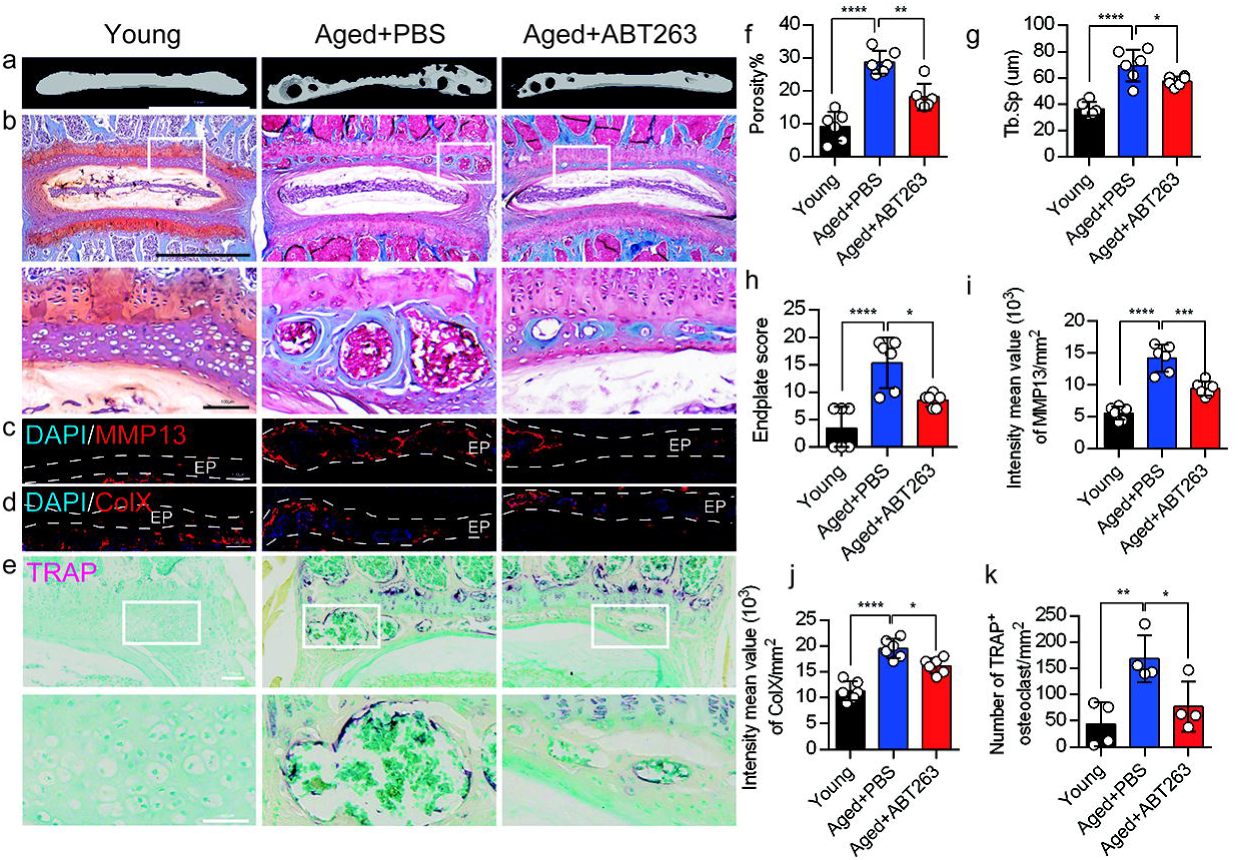
Depletion of SnOCs reduces spinal degeneration and sustains endplate microarchitecture in aged mice. (**a)** μCT images of the aged mouse caudal endplates of L4-L5 injected with PBS or ABT263. Scale bar, 1 mm. (**b**) Representative images of safranin O and fast green staining of coronal sections of the caudal endplates of L4-5 in aged mice caudal endplates of L4-L5 injected with PBS or ABT263, respectively. Lower panners are zoomed in images from upper white boxes. Scale bar, 1 mm (upper panels) and 100 μm (lower panels). (**c**) Representative images of spine degeneration marker MMP13 (red) and nuclei (DAPI; blue) staining in aged mouse caudal endplates of L4-L5 injected with PBS or ABT263. Scale bar, 100 μm. (**d**) Representative images of spine degeneration marker ColX (red) and nuclei (DAPI; blue) staining in aged mouse caudal endplates of L4-L5 injected with PBS or ABT263. Scale bar, 100 μm. (**e**) Representative images of TRAP (magenta) staining of coronal sections of the caudal endplates of L4-5 in aged mice caudal endplates of L4-L5 injected with PBS or ABT263, respectively. Lower panners are zoomed-in images from upper white boxes. Scale bar, 100 μm. (**f**) The quantitative analysis of the porosity percentage. (**g**) The quantitative analysis of the trabecular separation. (**h**) The endplate score based on the safranin O and fast green staining. (**i**) Quantitative analysis of the intensity mean value of MMP13 in endplates per mm^2^. (**j**) Quantitative analysis of the intensity mean value of ColX in endplates per mm^2^. (**k**) The quantitative analysis of the number of TRAP-positive cells in the endplate per mm^2^. *n* ≥ 3 per group. Statistical significance was determined by one-way ANOVA, and all data are shown as means ± standard deviations.

We next conducted μCT analysis and immunostaining in sham and LSI mice treated with ABT263 or PBS (Figure 5a-e) to determine the effect of SnOCs on endplate architecture, degeneration, and osteoclast formation in the context of spinal pain. In the LSI mice, ABT263 treatment significantly mitigated the porosity and trabecular separation of the caudal endplates of L4/5 compared to the PBS-treated group (Figure 5a, 5f, 5g). Safranin O staining and immunofluorescent staining of ColX and MMP13 within the endplate demonstrated ABT263 administration significantly reduced endplate score (Figure 5b, 5h), the distribution of MMP13 (Figure. 5c, 5i) and ColX (Figure 5d, 5j) and TRAP^+^ osteoclasts (Figure 5e, 5k) in the endplates compared to LSI mice treated with PBS. These findings collectively suggest the pivotal role of ABT263 in mitigating spinal degeneration and maintaining endplate remodeling in the context of spine pain.

**Figure 5.**
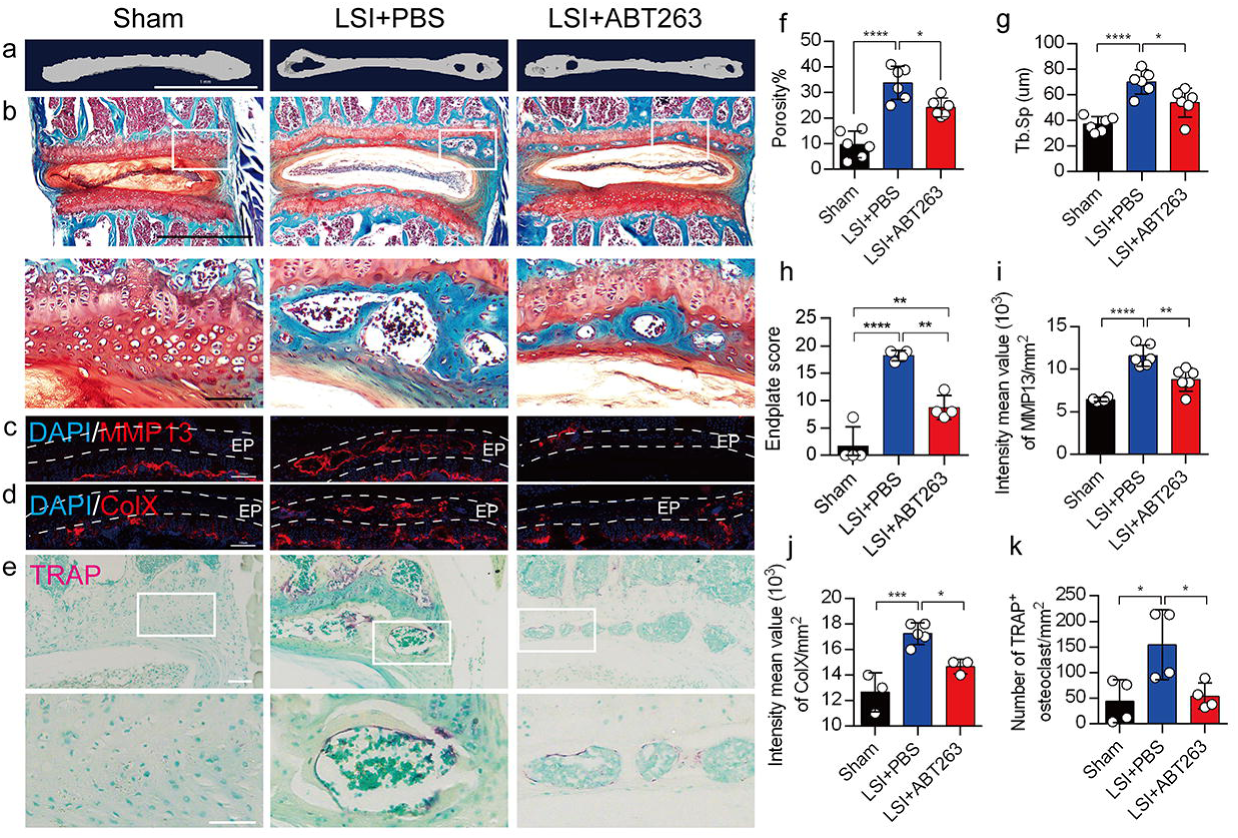
Depletion of SnOCs reduces spinal degeneration and sustains endplate microarchitecture in LSI mice. (**a)** μCT images of adult sham mice and 3-month-old LSI model mice caudal endplates of L4-L5 injected with PBS or ABT263. Scale bar, 1 mm. (**b**) Representative images of safranin O and fast green staining in different groups. Lower panners are zoomed-in images from upper white boxes. Scale bar, 1 mm (upper panels) and 100 μm (lower panels). (**c**) Representative images of immunofluorescent staining of spine degeneration marker MMP13 (red) and nuclei (DAPI; blue). Scale bar, 100 μm. (**d**) Representative images of immunofluorescent staining of spine degeneration marker ColX (red) and nuclei (DAPI; blue). Scale bar, 100 μm. (**e**) Representative images of TRAP (magenta) staining in different groups. Lower panners are zoomed-in images from upper white boxes. Scale bar, 100 μm. (**f**) The quantitative analysis of the porosity percentage of the mouse caudal endplates of L4-5 measured by the μCT. (**g**) The quantitative analysis of the trabecular separation (Tb.Sp) of the mouse caudal endplates of L4-5 measured by the μCT. (**h**) The endplate score based on the safranin O and fast green staining. (**i**) Quantitative analysis of the intensity mean value of MMP13 in endplates per mm^2^. (**j**) Quantitative analysis of the intensity mean value of ColX in endplates per mm^2^. (**k**) The quantitative analysis of the number of TRAP-positive cells in the endplate per mm^2^. *n* ≥ 3 per group. Statistical significance was determined by one-way ANOVA, and all data are shown as means ± standard deviations.

### Depletion of SnOCs abrogates sensory innervation and pain

We previously found that sensory innervation occurs in the porous endplates, contributing to spinal hypersensitivity, in LSI and aged mice^31^. To evaluate the contribution of SnOCs to sensory innervation and pain in LSI and aged mice, we co-stained for calcitonin gene-related peptide (CGRP), a marker of peptidergic nociceptive C nerve fibers, and PGP9.5, a broad marker of nerve fibers, in the endplates of LSI and aged mice treated with ABT263 or PBS (Figure 6a). Notably, we found fewer CGRP^+^ PGP9.5^+^ nerves in LSI mice and aged mice treated with ABT263 compared to those treated with PBS (Figure 6b-6e).

**Figure 6.**
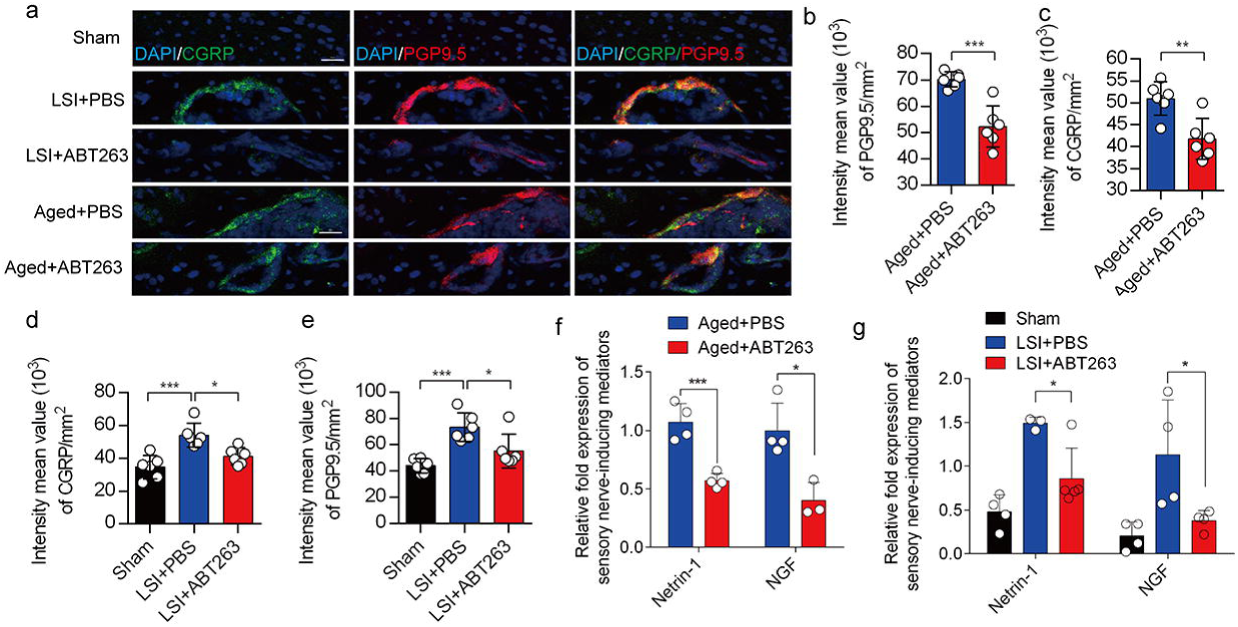
Depletion of SnOCs abrogates sensory innervation and pain in aged and LSI mouse models. (**a)** Representative images of immunofluorescent analysis of CGRP (green), PGP9.5 (red) and nuclei (DAPI; blue) of adult sham, LSI and aged mice injected with PBS or ABT263. Scale bar, 20 μm. (**b)** Quantitative analysis of the intensity mean value of PGP9.5 per mm^2^ in aged mice. (**c)** Quantitative analysis of the intensity mean value of CGRP per mm^2^ in aged mice. (**d)** Quantitative analysis of the intensity mean value of PGP9.5 per mm^2^ in the LSI mouse model. (**e)** Quantitative analysis of the intensity mean value of CGRP per mm^2^ in the LSI mouse model. (**f, g**) Relative fold expression of *Ntn* and *Ngf* in aged mice (**f**) or LSI mice (**g**) with or without ABT263 treatment. Statistical significance in panels b, c, and f are analyzed using *t*-tests, while panels d, e, and g are subjected to one-way ANOVA. All data are shown as means ± standard deviations.

To investigate the mechanisms underlying sensory innervation-induced spinal pain, we performed RT-qPCR to screen for expression of mediators regulating nerve fiber innervation and outgrowth, including Netrin-1 and NGF^42–44^. Compared to the young sham mice, aging and LSI was associated with significantly increased expression of *Ntn* and *Ngf* (the genes encoding netrin-1 and Ngf, respectively), which was substantially attenuated by ABT263 treatment in both LSI and aged mice (Figure 6f, 6g, Figure 6-figure supplement 2a, 2b).

Our earlier data showed that CGRP^+^ nociceptive nerve fibers and blood vessels were increased in the cavities of sclerotic endplates in the LSI and aged mice^31^. To study whether elimination of SnOCs prevent such blood vessel growth into the endplates, we co-stained for CD31, an angiogenesis marker (green), and Emcn, an endothelial cells marker (red), in the endplates of sham, LSI and aged mice treated with PBS or ABT263 (Figure 7a). In conjunction with the sensory nerve distribution within the porous endplates, we found noticeable growth of CD31^+^Emcn^+^ blood vessels into the endplates of the LSI and aged mice compared to young sham mice. This observation points toward an ongoing process of active ossification in the endplate. ABT263 treatment in the LSI and aged mouse models significantly mitigated the aberrant innervation of sensory nerves and blood vessels within the endplate compared to the PBS-treated mice (Figure 7b-7e). Collectively, these findings underscore that ABT263 treatment effectively reduces spinal hypersensitivity by diminishing the innervation of sensory nerves and blood vessels within the endplate.

**Figure 7.**
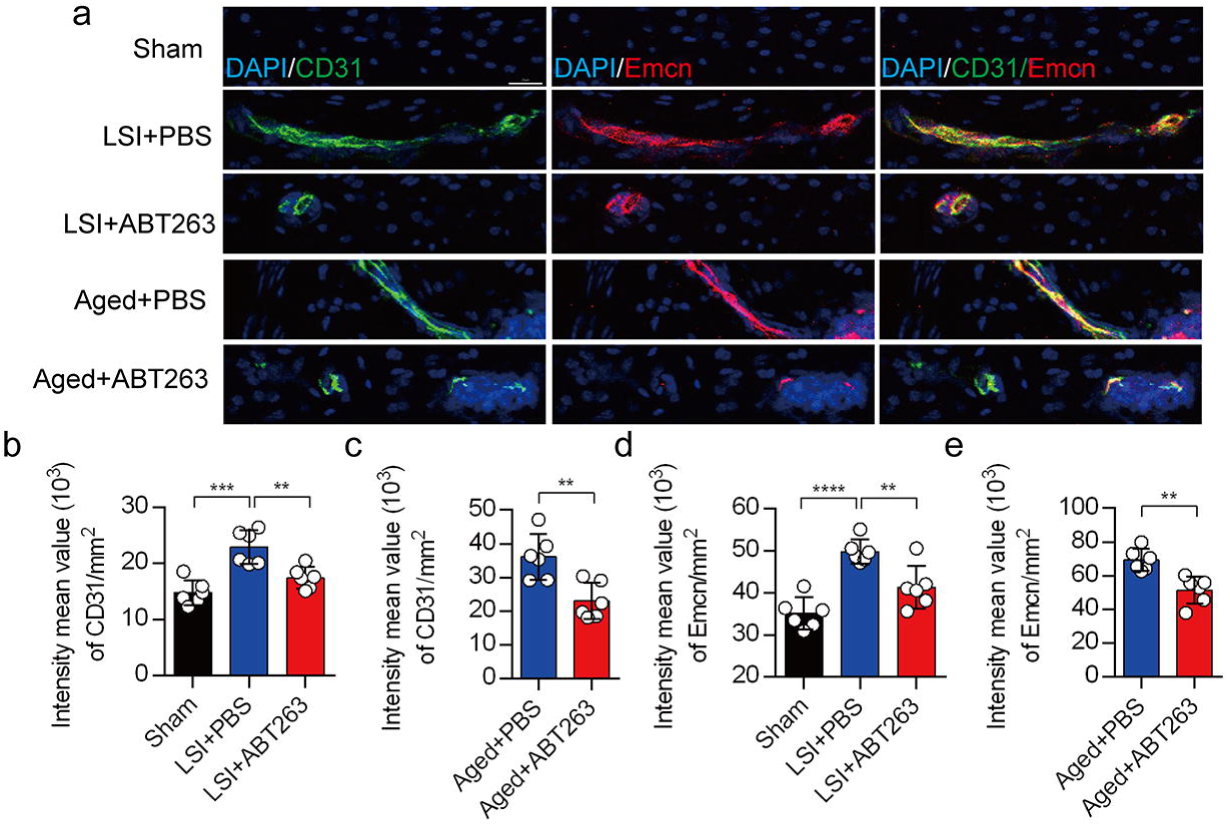
Depletion of SnOCs abrogates blood vessels innervation in aged and LSI mouse models. (**a)** Representative images of immunofluorescent analysis of CD31, an angiogenesis marker (green), Emcn, an endothelial cell marker (red) and nuclei (DAPI; blue) of adult sham, LSI and aged mice injected with PBS or ABT263. Scale bar, 20 μm. (**b)** Quantitative analysis of the intensity mean value of CD31 per mm^2^ in sham, LSI mice treated with PBS or ABT263. (**c)** Quantitative analysis of the intensity mean value of CD31 per mm^2^ in aged mice treated with PBS or ABT263. (**d)** Quantitative analysis of the intensity mean value of Emcn per mm^2^ in sham, LSI mice treated with PBS or ABT263. (**e)** Quantitative analysis of the intensity mean value of Emcn per mm^2^ in aged mice treated with PBS or ABT263. *n* ≥ 4 per group. Statistical significance was determined by one-way ANOVA, and all data are shown as means ± standard deviations.

## Discussion

LBP affects individuals of all ages and is a leading contributor to disease burden worldwide. Despite advancements in its assessment and treatment, the management of LBP remains a challenge for researchers and clinicians alike. Defects in a number of anatomical structures within the back may be responsible for back pain, including the intervertebral discs, facet joints, muscles, ligaments and nerve root sheaths. Of these, the intervertebral discs, facet joints and sacroiliac joints are implicated in the majority of the cases of LBP^40^. Furthermore, more than one structure may be contributing to the pain at any one time. During healing, neovascularisation occurs and minute sensory nerves can penetrate the disrupted annulus and nucleus pulposus, leading to mechanical and chemical sensitization^41^.

During LSI or aging, endplates undergo ossification, leading to elevated osteoclast activity and increased porosity^36,42–44^. The progressive porous transformation of endplates, accompanied by a narrowed intervertebral disc (IVD) space, is a hallmark of spinal degeneration^44,45^. Considering that pain arises from nociceptors, it is plausible that LBP may be attributed to sensory innervation within endplates. Additionally, porous endplates exhibit higher nerve density compared to normal endplates or degenerative nucleus pulposus^46^. Netrin-1, a crucial axon guidance factor facilitating nerve protrusion, has been implicated in this process^47–49^. The receptor mediating Netrin-1-induced neuronal sprouting, deleted in colorectal cancer (DCC), was found to co-localize with CGRP^+^ sensory nerve fibers in endplates after LSI surgery^50,51^. In summary, during LSI or aging, osteoclastic lineage cells secrete Netrin-1, inducing extrusion and innervation of CGRP^+^ sensory nerve fibers within the spaces created by osteoclast resorption. This Netrin-1/DCC-mediated pain signal is subsequently transmitted to the dorsal root ganglion (DRG) or higher brain levels. In our previous study, we found that osteoclasts induce sensory innervation of the porous areas of sclerotic endplates, which induced spinal hypersensitivity in LSI-injured mice and in aging. Inhibition of osteoclast formation by knockout of *Rankl* in the osteocytes significantly inhibits LSI-induced porosity of endplates, sensory innervation and spinal hypersensitivity^31^. Likewise, knockout of *Ntn1* in osteoclasts abrogates sensory innervation into porous endplates and spinal hypersensitivity^31^. In an osteoarthritis (OA) mouse model, we found a role for osteoclast-secreted netrin-1 in the induction of sensory nerve axonal growth in the subchondral bone. Reduction of osteoclast formation by knockout of *Rankl* in osteocytes inhibited the growth of sensory nerves into subchondral bone, DRG neuron hyperexcitability and behavioral measures of pain hypersensitivity^52^. Our previous study revealed that osteoclast-lineage cells may promote both nerve and vessel growth in osteoarthritic subchondral bone, leading to disease progression and pain^53^. Here, we report that SnOCs are mainly responsible for modulating the secretion of netrin-1 and NGF, which mediate sensory innervation and induce hypersensitivity of spine.

Osteoclasts are the principal bone-resorbing cells essential for bone remodeling and skeletal development. Here, we report that osteoclasts in the endplate of the vertebral column undergo cellular senescence during injury and aging. The senescence process is programmed by a conserved mechanism because it is restricted to a specific region and follows a specific time course. Cellular senescence was defined by the presence of a senescence marker, SA-βGal, and a key senescence mediator, p16INK4a, detected in the bone tissue sections. In the present study, we found that the number of TRAP^+^ and SA-βGal^+^ or p16^+^ senescent osteoclasts in endplates was significantly increased in LSI injured mice and aged mice compared to sham injured mice with PBS treatment. These findings support our hypothesis that increased numbers of SnOCs in LSI or aging conditions contribute to nerve innervation factors secretion, which leads to spine pain.

Current LBP management strategies have limited therapeutic effects, and progressive pathological spinal changes are observed frequently with these treatments. According to the American College of Physicians guidelines, pharmacological recommendations for acute or subacute LBP should begin with NSAIDs or muscle relaxants (moderate-quality evidence). There is no consensus on the duration of NSAID use, and caution is advised with persistent use due to concerns for cardiovascular and gastrointestinal adverse events. Guidelines by the American College of Physicians^8^ recommend tramadol or duloxetine as a second-line treatment and opioids as the last-line treatment for chronic LBP. A meta-analysis showed that opioids offer only modest, short-term pain relief in patients with chronic LBP^54^. The addictive potential of opioids coupled with several side effects limits their use in the management of such pain^8^. Consequently, these drugs provide insufficient and unsustained pain relief with considerable adverse effects.

A clinical study demonstrated that nerve density is higher in porous endplates than in normal endplates and is associated with pain. Radiofrequency denervation treatment can be used for pain relief originating from the lumbar facet joints^55,56^. However, National Institute for Health and Care Excellence (NICE) guidelines from the United Kingdom^57^ recommend considering radiofrequency denervation only when the main source of pain originates from the facet joints, when pain is moderate to severe, and only when evidence-based multidisciplinary treatment has failed. During LBP progression, sensory nerves and blood vessels are aberrantly innervated in the endplate, which leads to pain. In this study, we aimed to reduce pain by decreasing nerve innervation. We found that SnOCs mediate nerve fiber innervation by elevated secretion of netrin-1and NGF. Moreover, our previously study reported that osteoclasts can secret netrin-1 to attract sensory nerve growth^31^. NGF can be produced by osteoblasts in response to mechanical load^58^ or by bone marrow stromal cells (BMSCs), and Sema3A is secreted by osteoblasts^59^. We believe a cross-talk exists between osteoclasts and osteoblasts or BMSCs after LSI or aging to modulate the secretion of nerve fiber innervation mediators.

The recent discovery that senescent cells play a causative role in aging and in many age-related diseases suggests that cellular senescence is a fundamental mechanism of aging. In aging, as well as in metabolic disorders, the immune response is affected by senescent cells that no longer replicate, but have a senescence-associated secretory phenotype that produces high levels of proinflammatory molecules. Certain chemotherapy drugs, known as senolytics, however, kill senescent cells and other drugs, called anti-SASP drugs, block their pro-inflammatory cell signaling. ABT263 is a selective BCL-2 and BCL-xL inhibitor and is one of the most potent and broad-spectrum senolytic drugs. In the current study, we explored the role of ABT263 to manage spinal pain, and we found to it could effectively clear SnOCs cells in two mouse models. A decreased number of SnOCs will relieve pain by decreasing sensory neuron innervation in the endplate. However, ABT263 does not specifically eliminate SnOCs and thus further studies are required to prove the role of SnOCs in spinal hypersensitivity. Furthermore, ABT263 usually possesses various on-target and/or off-target toxicities, which could preclude its clinical use. Even so, for the first time to the best of our knowledge, we show evidence that SnOCs promote LBP by neurotrophic-mediated pathways. Our findings suggest that depletion of SnOCs, perhaps by use of a senolytic, can reduce sensory innervation and attenuate LBP, thus representing a new avenue in the management of this widespread condition (Figure 8).

**Figure 8.**
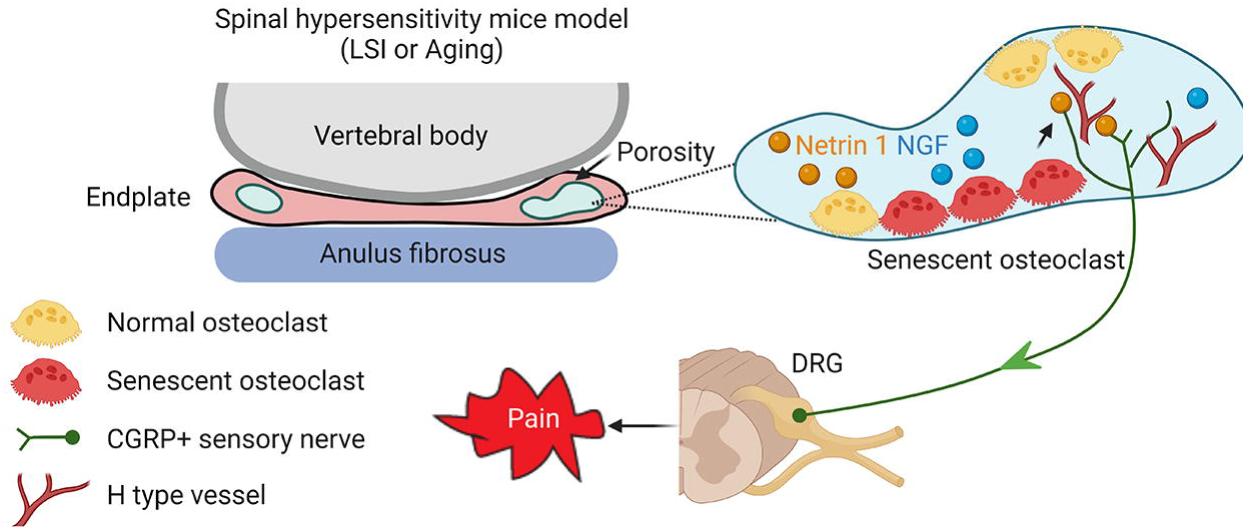
Schematic diagram of SnOCs in porous endplate-induced spinal pain. In LSI or aging mouse models there is an induction of spinal hypersensitivity due to increased numbers of SnOCs in the endplate, leading to excessive secretion of sensory nerve mediators, such as Netrin 1, to attract CGRP^+^ sensory nerve innervation. Additionally, aberrant microarchitecture and spinal degeneration are associated with increased wiring of sensory nerve fibers and H-type vessels in the endplate, all further contributing to lower back pain.

## Methods

### Mice

Three- and 24-month-old C57BL/6J male mice were purchased from Jackson Laboratory. Three-month-old mice were anesthetized with ketamine (Ketalar, 0.13 mg/kg, intraperitoneally) and xylazine (Millipore sigma; PHR3264, 12 mg/kg, intraperitoneally). Then, the L3–L5 spinous processes, and the supraspinous and interspinous ligaments were resected to induce instability of the lumbar spine and to create the LSI model^36–38^. For mice in the sham group, we only surgically detached the posterior paravertebral muscles from L3-L5. ABT263 (Navitoclax, Selleckchem, S1001) was administered to mice by gavage at 50 mg/kg per day for 7 days per cycle for two cycles with a 2-week interval between the cycles (the whole treatment time was 4 weeks). ABT263 was administered to aged (24 M) C57BL/6J mice (12 per group) at the age of 23 months. In the meantime, 4-month-old C57BL/6J mice 4-weeks post LSI or sham-operation were treated with ABT263 or vehicle (PBS)^39^. The control group (sham group) for the LSI group refers to C57BL/6J mice that did not undergo LSI surgery, while the control group (young group) for the aged group refers to 4-month-old C57BL/6J mice. All mice were maintained at the animal facility of The Johns Hopkins University School of Medicine (protocol number: MO21M276, MO21M270, MO22M18). All experimental protocols were approved by the Animal Care and Use Committee of The Johns Hopkins University, Baltimore, MD.

### Behavioral testing

Behavioral tests were performed after ABT263 administration and before sacrifice. All behavioral tests were performed by the same investigator, who was blinded to the study groups. The hind paw withdrawal frequency in response to a mechanical stimulus was determined using von Frey filaments of 0.07 g and 0.4 g (Aesthesio® Precision Tactile Sensory Evaluator). Mice were placed on a wire metal mesh grid covered with a clear plastic cage. Acclimatization of the animals in the enclosure was for 30 min. We applied von Frey filaments to the mid-plantar surface of the hind paw through the mesh floor with enough pressure to buckle the filaments. A trial consisted of 10 times at 1-s intervals. Mechanical withdrawal frequency was calculated as the number of withdrawal times in response to ten applications after three replicates.

The Hargreaves Test of nociception threshold was evaluated by the Model Heated 400 Base. The animals were transferred from the holding room to the enclosure and acclimatization of the animals in the enclosure was for 30 min. The duration of exposure before the hind paw withdrawn with three replicates is recorded after focusing the mouse’s mid-plantar surface of the hind paw with the light.

Spontaneous wheel-running activity was recorded using activity wheels designed for mice (model BIO-ACTIVW-M, Bioseb)^60^. The software enabled recording of activity in a cage similar to the mice’s home cage, with the wheel spun in both directions. The device was connected to an analyzer that automatically recorded the spontaneous activity. We evaluated the distance traveled, maximum speed and total active time during 2 days for each mouse.

### RT-qPCR

Mice were euthanized with an overdose of isoflurane inhalation. Total RNA was then extracted from the L4–L5 lumbar spine endplate tissue samples. Briefly, spine endplate tissue was ground in liquid nitrogen and isolated from Buffer RLT Plus using RNeasy Plus Mini Kits (Qiagen, Germany) and homogenized directly with ultrasound (probe sonication at 50 Hz for three times, 10 s per cycle). The purity of RNA was measured by the absorbance at 260/280 nm. Then, the RNA was reverse transcribed into complementary DNA by PrimeScript RT (reverse transcriptase) using a primeScript RT-PCR kit (Takara), and qPCR was performed with SYBR Green-Master Mix (ThermoFisher Scientific, USA) on a QuantStudio 3 (Applied Biosystems, USA). Relative expression was calculated for each gene by the 2-^△△^CT method, with glyceraldehyde 3-phosphate dehydrogenase (*GAPDH*) used for normalization. Primers used for RT-qPCR are listed below: *Netrin 1*: forward: 5’-CCTGTCACCTCTGCAACTCT-3’, reverse: 5’-TGTGCGGGTTATTGAGGTCG-3’; *NGF*: forward: 5’-CTGGCCACACTGAGGTGCAT-3’, reverse: 5’-TCCTGCAGGGACATTGCTCTC-3’; *BDNF*: forward:5’-TGCAGGGGC ATAGACAAAAGG-3’, reverse: 5’-CTTATGAATCGCCAGCCAATTCTC-3’; *NT3*: forward: 5’-CTCATTATCAAGTTGATCCA-3’, reverse: 5’-CCTCCGTGGTGATGTTCTATT-3’; *Slit3*: forward: 5’-AGT TGTCTGCCTTCCGACAG-3’, reverse: 5’-TTTCCATGGAGG GTCAGCAC-3’; *GAPDH*: forward: 5’-ATGTGTCCGTCGTGGATCTGA-3’, reverse: 5’-ATGCCTGCTTCACCACCTTCTT-3’.

### ELISA (enzyme-linked immunosorbent assay)

We determined the concentration of Netrin 1 (LSBio, LS-F5882) and NGF (Boster, EK0470) in the L3 to L5 endplates using the ELISA Development Kit according to the manufacturer’s instructions.

### μCT

Mice were euthanized with an overdose of isoflurane inhalation and flushed with PBS for 5 min followed by 10% buffered formalin perfusion for 5 min via the left ventricle. Then, the whole lumbar spine was dissected and fixed in 10% buffered formalin for 48 h, transferred into PBS, and examined by high-resolution μCT (Skyscan1172). The scanner was set at a voltage of 55LkV, a current of 181 μA and a resolution of 9.0 μm per pixel to measure the endplates and vertebrae. The ribs on the lower thoracic spine were included for identification of L4–L5 unit localization. Images were reconstructed and analyzed using NRecon v1.6 and CTAn v1.9 (Skyscan US, San Jose, CA), respectively. Coronal images of the L4–L5 unit were used to perform three-dimensional histomorphometric analyses of the caudal endplate. The three-dimensional structural parameters analyzed were total porosity and trabecular bone separation distribution (Tb.Sp) for the endplates. Six consecutive coronal-oriented images were used for showing 3-dimensional reconstruction of the endplates and the vertebrae using three-dimensional model visualization software, CTVol v2.0 (Skyscan US).

### Histochemistry, immunohistochemistry and histomorphometry

After μCT scanning, the spine samples were decalcified in 0.5 M EDTA (pH 7.4) for 30 days and embedded in paraffin or optimal cutting temperature compound (Sakura Finetek, Torrance, CA).

Four-μm-thick coronal-oriented sections of the L4–L5 lumbar spine were processed for Safranin O (Sigma–Aldrich, S2255) and fast green (Sigma–Aldrich, F7252) staining, TRAP (Sigma– Aldrich, 387A-1KT) staining, and immunohistochemistry staining with an established protocol^31^. Thirty-μm-thick coronal-oriented sections were prepared for blood vessel-related immunofluorescent staining, and 10-µm-thick coronal-oriented sections were used for other immunofluorescent staining.

The sections were incubated with primary antibodies to mouse Col X (1:100, ab260040, Abcam), MMP13 (1:100, ab219620, Abcam), Endomucin (1:100, sc-65495, Santa Cruz Biotechnology), CD31 (1:100, 550389, BD Biosciences), CGRP (1:100, ab81887, Abcam), PGP9.5 (1:100, SAB4503057, Sigma-Aldrich), Netrin-1 (1:100, ab39370, Abcam), TRAP (1:100, PA5-116970, invitrogen), overnight at 4 °C. Then, the corresponding secondary antibodies and 4′,6-diamidino-2-phenylindole (DAPI, Vector, H-1200) were added onto the sections for 1 h while avoiding light.

The sample images were observed and captured by the confocal microscope (Zeiss LSM 780). Image J (NIH) software was used for quantitative analysis. We calculated endplate scores as described previously^61,62^.

### Statistics

All data analyses were performed using SPSS, version 15.0, software (IBM Corp.). Data are presented as meansL±Lstandard deviations. Unpaired, two-tailed Student’s *t*-tests were used for comparisons between two groups. One-way ANOVA with Bonferroni’s post hoc test was used for comparisons among multiple groups. For all experiments, pL<L0.05 was considered to be significant. There were no samples or animals that were excluded from the analysis. The experiments were randomized, and the investigators were blinded to allocation during experiments and outcome assessment.

## Acknowledgments

This research was supported by the United States NIH National Institute on Aging under award numbers R01AG068997, P01AG066603, R01AG076783, R01AR071432 (to X.C.).

## Figure legends

**Figure 1-figure supplement 1.** (**a)** Representative images of immunofluorescent analysis of HMGB1, a senescent marker (green), Trap, an osteoclast marker (red) and nuclei (DAPI; blue) of adult sham, LSI and aged mice. (**b)** Quantitative analysis of the number of Trap^+^HMGB1^+^ SnOCs per mm^2^. *n* ≥ 4 per group. Statistical significance was determined by one-way ANOVA, and all data are shown as means ± standard deviations.

**Figure 6-figure supplement 1.** (**a)** ELISA analysis showing the concentration of Netrin1 in L3-5 endplates of adult sham, LSI+PBS and LSI+ABT263 mice. (**b)** ELISA analysis showing the concentration of NGF in L3-5 endplates of adult sham, LSI+PBS and LSI+ABT263 mice. *n* = 3 per group. Statistical significance was determined by one-way ANOVA, and all data are shown as means ± standard deviations.

## Notes

### Competing Interest Statement

The authors have declared no competing interest.

### Summary of Updates

A white rectangle (high magnification frame) in Figure 4b (Aged+ABT263 group) was added; The staining labeling in Figure 2b was updated; The animal protocol number was updated. All of these revisions are consistent with the final published version.

