## Supplementary figures and images for "Senescence of endplate osteoclasts induces sensory innervation and spinal pain"

### Figure 1-figure supplement 1

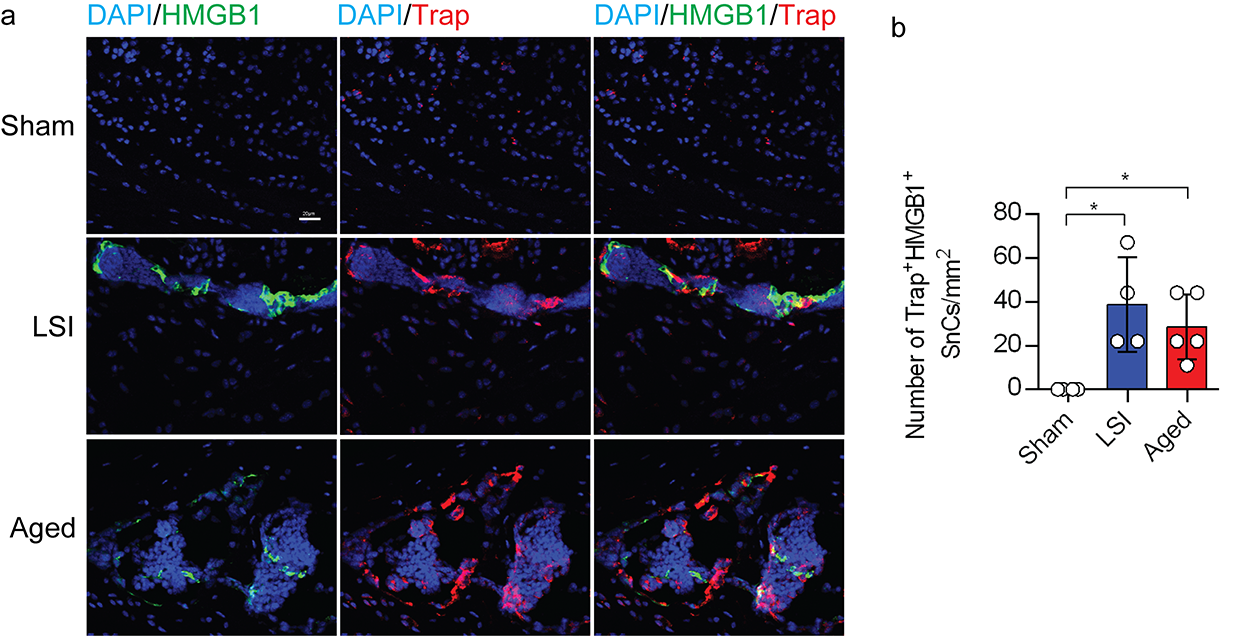

### Figure 6-figure supplement 1

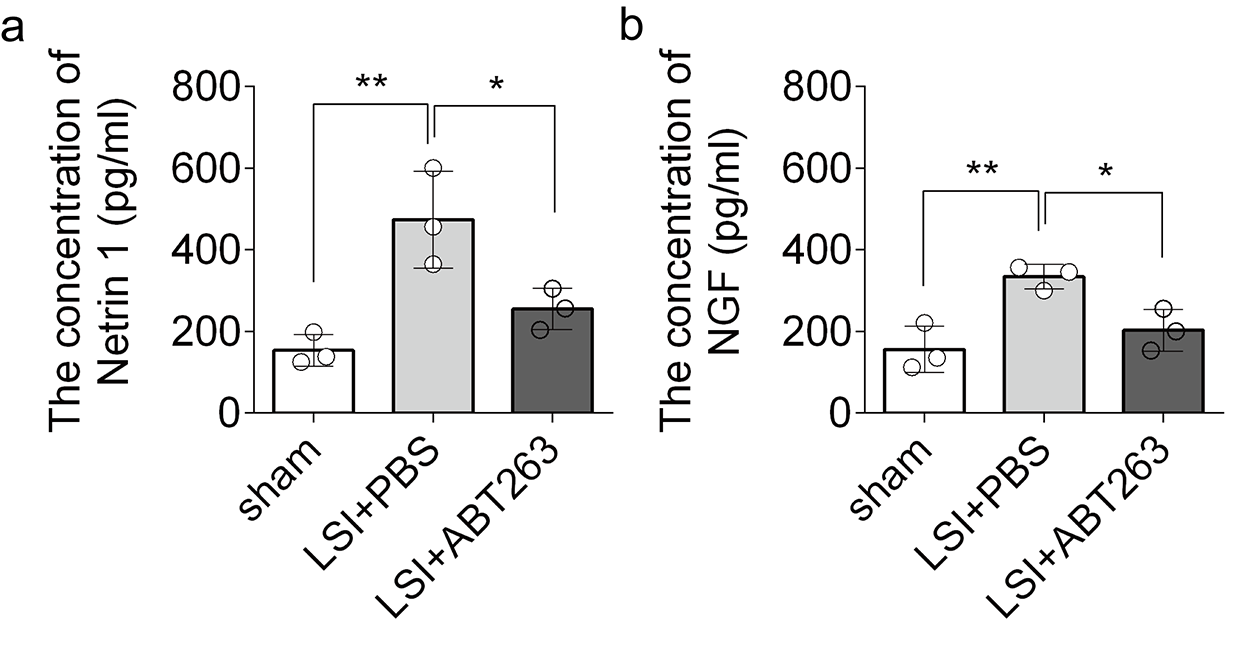
